# Lexical brain responses in 10-year-old children are impaired in dyslexia: an FPVS-EEG study

**DOI:** 10.1101/2025.07.20.665366

**Authors:** Claire Gigleux, Alice van de Walle de Ghelcke, Christine Schiltz, Bruno Rossion, Aliette Lochy

**Affiliations:** Department of Behavioural and Cognitive Sciences, Faculty of Humanities, Social and Educational Sciences, Institute of Cognitive Science and Assessment, University of Luxembourg, Esch-sur-Alzette, Luxembourg; Psychological Sciences Research Institute (IPSY), Université Catholique de Louvain, Louvain-la-Neuve, Belgium; Université de Lorraine, CNRS, IMoPA, F-54000, Nancy, France; Université de Lorraine, CHRU-Nancy, Service de Neurologie, Nancy, France

## Abstract

The developmental origin of the left occipitotemporal cortex specialization for automatic lexical access from vision remains unclear. Here we investigated cortical specialization for print processing in children with or without dyslexia, focusing on two distinct tuning levels: coarse-grained tuning for letter/symbol discrimination, and fine-grained tuning for word/pseudoword discrimination. 10-year-old typical readers (n=24) and children with dyslexia (n=14) were tested with electroencephalography (EEG) and fast periodic visual stimulation (FPVS), viewing streams of stimuli at a relatively fast rate (6Hz) for 40 seconds with deviant categories every 5 items (at 6Hz/5=1.2Hz). Deviant words or pseudowords among pseudo-font strings elicited clear coarse ocicpito-temporal discrimination responses significantly larger over the left than the right hemisphere (LH), numerically larger in typical readers. Unlike in adults, these responses were unaffected by lexicality. Deviant regular or irregular words among matched pseudowords generated a finer-grained word-selective response only over the LH. While irregular words elicited similar brain responses in both groups, regular words were not discriminated from pseudowords in children with dyslexia. These results demonstrate the sensitivity of FPVS-EEG to implicitly detect lexical neural responses in 10 years old children within a few minutes, as well as atypical lexical processing in children with dyslexia.

## INTRODUCTION

In alphabetic writing systems, the acquisition of the orthographic code is a multi-stage process that begins in early childhood and entails mastering letter-to-sound correspondences. In the left hemisphere, the ventral occipitotemporal cortex (VOTC) specializes to process written words (Dehaene-Lambertz et al. 2018; Wandell et al., 2012). During literacy acquisition, this region is functionally segregated to handle different aspects of visual word recognition in its posterior part (Lerma-Usabiaga et al., 2018; Caffarra et al., 2021). Visual word processing can be conceptualized as occurring along a continuum, from coarse to fine levels of visual processing. Coarse tuning enables the differentiation between orthographic and non-orthographic stimuli such as symbols or false fonts. Fine tuning refers to the brain’s capacity to discriminate between orthographic stimuli, including non-words (unpronounceable letter strings), pseudo-words (pronounceable letter strings), and frequent words. The present study aims to provide implicit neural markers of these two levels of print processing in ten-year-old children with or without reading disorder.

### Developmental emergence of neural sensitivity to print

The coarse-grained level of orthographic processing illustrates the emergence of a broad sensitivity to print pattern. Most kindergarten children already know some letters by the age of 5. They also demonstrate knowledge of visual word features by manipulating word length, letter type, and digrams to imitate their native spelling (Treiman et al., 2018), suggesting the development of general awareness to print. At the neural level, print expertise has been assessed in ERP and fMRI studies by comparing responses to print vs. an unrelated baseline condition such as symbols (Maurer et al., 2006; Araújo et al., 2015) or false fonts (Maurer et al., 2005, Olulade et al., 2013; Centanni et al., 2017). With EEG, the N170 displays larger amplitudes for print over the left hemisphere (Maurer et al., 2005) and typically emerges only after 1 to 1.5 years of formal education (Zhao et al., 2014), although it has also been measured earlier in specific training designs (Brem et al., 2010). However more recently, fMRI studies showed greater activation for print immediately after the beginning of formal literacy (Dehaene-Lambertz et al., 2018). Also, studies relying on an alternative frequency-tagging approach using Fast Periodic Visual Stimulation (FPVS) with EEG recordings and an oddball design have revealed reliable left occipito-temporal cortex discrimination of real letter strings from pseudo-letters in pre-reading kindergartners (Lochy et al., 2016; van de Walle de Ghelcke et al., 2021). This suggests that coarse-grained processing (letters > symbols) may be established prior to any explicit reading instruction. This fits with behavioral studies showing that upper- and lower-case letters of the alphabet are recognized as letters at that age, this ability furthermore laying the foundation for later letter-sound mapping (Paige et al., 2018).

As reading experience progresses during development, the system refines its tuning to visual print by increasing its sensitivity to different forms of letter strings, such as words, pseudowords, and non-words. From 7 years of age, typically developing children have knowledge of the orthographic rules of their writing system, which allows them to identify permissible letter combinations and positions (Pacton et al., 2001; 2002). Greater N170 amplitude for legal (words, pseudo-words) over illegal (non-words) letter strings has been found in studies with 10-year-old children (Coch & Mitra, 2010; Coch & Meade, 2016) and even in proficient 7 years-old German and Chinese readers (Tong et al., 2016; Zhao et al., 2014). However, at a finer level of visual print tuning, the specialization within the category of orthographically legal strings for real words over word-like stimuli (pseudowords) requires complementary investigation. A systematic review by Amora et al. (2022) examined the lexicality effect (words > pseudo-words) across alphabetic and logographic languages in school-aged children (7 to 11 years-old). In this review, a reverse lexicality effect i.e., more negative N170 for pseudowords, was reported in older age groups of Chinese readers (9 to 11 years old) as found by Zhao et al. (2019). In contrast, three studies found no lexicality effect: Eberhard-Moscicka et al. (2015) in German readers, Tong et al. (2016) in Chinese readers, and Zhao et al. (2019) in their younger age groups of Chinese readers. In the study by Coch & Meade (2016) comparing 3^rd^, 4^th^ and 5^th^ graders, the amplitude of the N170 did not differ between words and pseudowords in any group, but the N1 latency was shorter for words than pseudowords, in 4^th^ graders only (not anymore in 5^th^ graders). This lack of consistency across developmental studies could well reflect the lack of sensitivity of the N170 to fine-grained discrimination, as there are contradictions even in adulthood (Wydell et al., 2003; Pammer et al., 2004). Thus, until now, there is no clear picture on the developmental emergence of brain specialization for words over pseudowords. It could be that such fine tuning to real words involves a long time course and protracted development (Coch & Meade, 2016), but it could also be a matter of the (lack of) sensitivity of the experimental approach. Therefore, we investigated this issue with FPVS-EEG in 4^th^ and 5^th^ graders given that behavioral studies suggest that fast lexical recognition of words typically develops around that age, as evidenced by reduced length effect and increased reading fluency (Ballot & Zesiger, 2024; Barton et al., 2014; Karageorgos et al., 2019)

### Word regularity

While literacy skills develop similarly across alphabetic writing systems (Ziegler et al., 2010; Landerl et al., 2022), language-specific orthographic characteristics influence pattern of reading performance and impairment (Ziegler et Goswami, 2005; Caravolas, 2018; Carioti et al., 2021; Reis et al., 2023). In French language, lexical responses may depend on the regularity of the words. This concept refers to how consistently a word adheres to the phonological rules that govern the conversion of letters (graphemes) into sounds (phonemes) in a language. Therefore, irregular words (e.g., “femme” read [fam], “gars” read [ɡɑ]) involve more complex cognitive processes, relying on vocabulary (Krepel et al., 2021), and orthographic knowledge (Nash et al., 2023) to overcome the challenges posed by their grapheme-phoneme correspondences. In the case of dyslexia, regular words may be harder to decode due to impaired phonological processes (Boets et al., 2013; Ramus & Szenkovits, 2008). Irregular words may also induce specific difficulties as they rely on lexical processes requiring more orthographic knowledge and vocabulary, that are also areas of weakness in dyslexia (Wang et al., 2013; Lovett, 1994). It is therefore of interest to assess whether automatic French word recognition can be modulated by orthographic regularity, across reading abilities.

### The case of Dyslexia

Dyslexia is a developmental reading disorder (DSM-5) characterized by significant and persistent difficulties with accurate and fluent word recognition, which complexifies orthographic processing both at coarse- and fine-grained levels. Indeed, the reading network of individuals with dyslexia shows a distinct developmental trajectory, from pre-reading stages to later reading difficulties (Chyl et al., 2021). Neuroimaging studies of individuals with dyslexia have consistently shown reduced activation of various VOTC areas (Pina Rodrigues et al., 2019; Danelli et al., 2017; Debska et al., 2021). As a consequence, the disorder has been associated with reduced print sensitivity (Maurer et al., 2007), deficits in orthographic and phonological processing (Araujo et al., 2012, 2015; Mahé et al., 2018) and impaired access to the lexicon (Schulte-Körne et al., 2004). Over-activation in the left inferior frontal cortex and other regions during reading suggest compensatory mechanisms (Paulesu et al., 2014; Cainelli et al., 2023). In EEG studies, N170 amplitude differences for coarse-tuning in individuals with dyslexia were not observed for letter vs. non-letter strings across orthographies (9-13 year old: Araujo et al., 2012; 6-9 year old Maurer et al., 2007), or were found to be reduced (23 year old: Araujo et al., 2015) or absent in fine-tuning for words vs. pseudowords (20-27 year old: Shaul, 2013; 16-17 year old: Taroyan et al., 2009). Notably, these differences in neural processes are thought to be particularly pronounced between the ages of 6 and 8, with some normalization by the age of 12, although differences in literacy skills remain significant (Chyl et al., 2021; Morken et al., 2017). In this context, the present study aims to investigate differences in visual word recognition at the coarse and fine levels in 10-year-old children with and without dyslexia.

### The current study

The main objectives of the present study are (1) to characterize the lexical neural responses of 4^th^ and 5^th^ graders (average 10-year-old children) at both coarse and fine levels of orthographic processing using FPVS and (2) to compare the pattern of neural responses in aged-matched children with dyslexia.

In FPVS, stimuli are presented at a relatively fast periodic rate (e.g., six stimuli per second) inducing an objectively defined neural response at the stimulation frequency. In the oddball FPVS paradigm used here, a base category (e.g., pseudo-fonts) is interspersed with a “deviant” or oddball stimulus category (e.g. a word) inserted at a slower periodic rate (e.g., 1.2 Hz or one oddball every five stimuli) (Heinrich et al., 2009; see Rossion et al., 2020 for review). The common neural response to all stimuli converges at the base frequency of presentation (e.g., 6 Hz), while responses specific to the oddball category are elicited at 1.2 Hz and its harmonics (Lochy & Schiltz, 2019; Rossion et al., 2020). The oddball response is indicative of the brain’s differential processing mechanisms for the contrasted categories of stimuli, such as false letter strings (base’) versus letter strings (‘oddball’) (Lochy et al., 2016). Thanks to its high signal-to-noise ratio (Regan, 1989; Norcia et al., 2015), this FPVS-EEG approach provides a unique window into the visual system’s specialization for processing letters.

This EEG-based method provides another tool for comparing lexical responses in typical readers and children with dyslexia without the need for an explicit linguistic task, thus avoiding the challenges arising from reading activities. In recent years, FPVS-EEG has demonstrated its sensitivity to differentiate between poor and proficient readers (Lochy et al., 2016; van de Walle de Ghelcke, 2020; Lutz et al., 2024; Marchive et al., 2025), manifesting as a reduction of response amplitudes in poor readers. Recently, this approach was used to assess lexical discrimination (words among pseudowords) in dyslexic adults (Lochy et al., 2025). Besides an overall reduced response amplitude to words in adults with dyslexia, results also showed a striking pattern in this population. Indeed, regular words did not trigger any discrimination response among pseudowords, while irregular words did. The authors interpreted this finding as revealing a contextual influence of pseudoword base stimuli, in a dual-route perspective: sequences presented streams of mostly pseudowords, which presumably tuned the reading system to sublexical decoding. Regular words, given that they may also be processed with this pathway, did not automatically activate lexical recognition. Irregular words, to the contrary, generated a discrimination response because they obligatorily activate lexical recognition processes. This finding thus suggests a possible modulation of lexical/non-lexical processes due to the contextual enhancement of the non-lexical route by pseudowords, and will be further tested in the current study.

Here in young children, we assessed two levels of discrimination. First, a coarse-grained level is assessed by means of oddball letter strings (words or pseudowords) embedded in pseudo-font base stimuli. Therefore, base–oddball stimuli differed in broad visual characteristics (**Figure 1**). Given their representation in the lexicon, we hypothesized that oddball words will elicit a more robust cortical response than oddball pseudowords and that this amplitude difference should be lower in children with dyslexia. Second, a fine-grained level is assessed by means of irregular or regular French words embedded in pseudoword base stimuli. Given that oddball words are presented in a stream of pseudoword sequences that presumably trigger decoding mechanisms, we hypothesized that irregular words, which can only be processed by lexical mechanisms, should elicit a larger amplitude response than regular words, which can be processed by the two routes (Coltheart et al., 2001). Furthermore, discrimination of words in pseudowords should be reduced in children with dyslexia, given that lexical representation benefits mainly readers with a certain level of proficiency. Finally, we also wanted to assess whether we could replicate the specific pattern observed in adults with dyslexia described above (Lochy et al., 2025). In that case, regular words should not be discriminated from pseudowords by dyslexic children only, revealing a lack of switch between decoding and lexical processes, while irregular words would be discriminated given that they must be processed by lexical mechanisms.

**Fig. 1.**
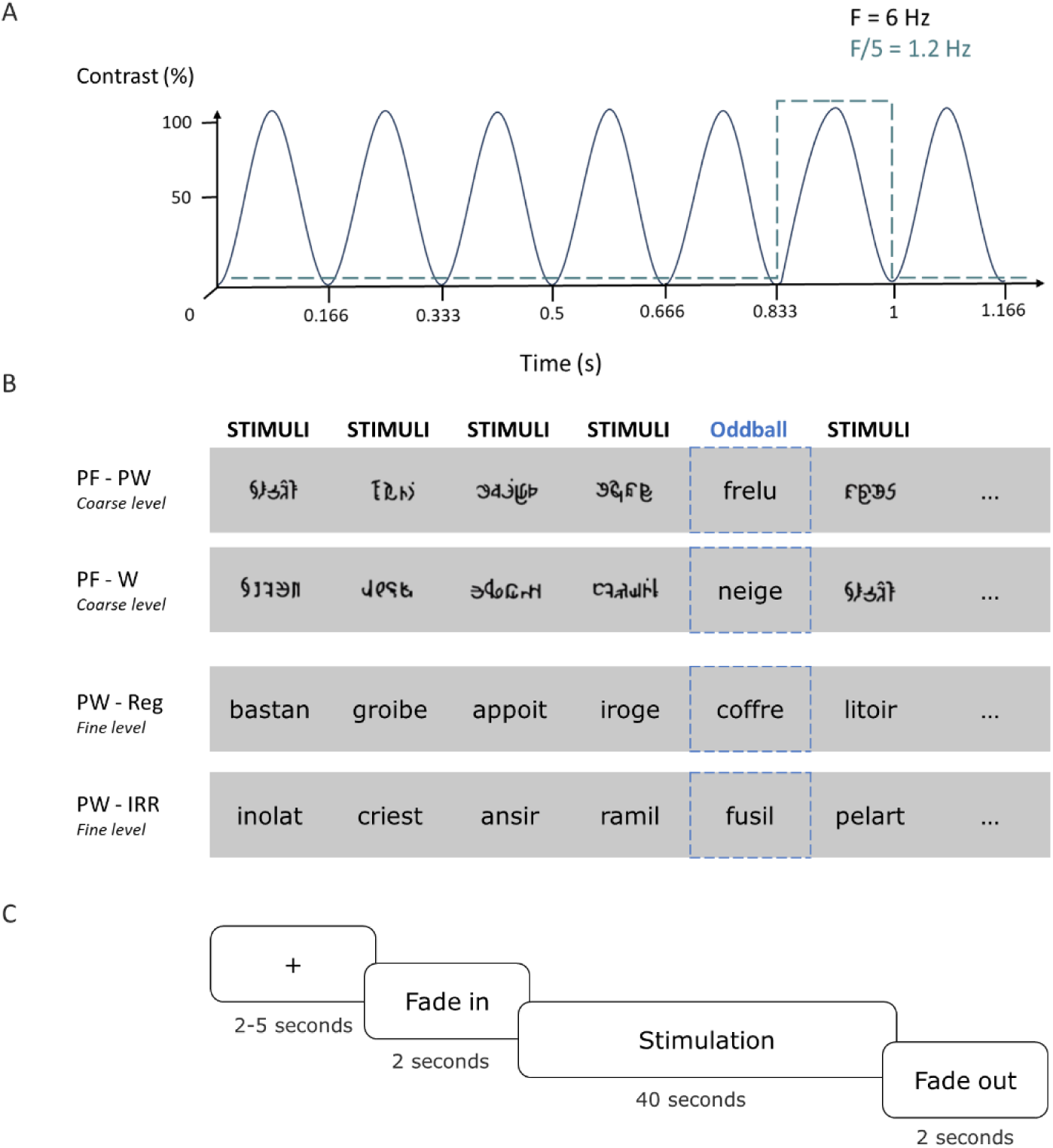
Experimental design. **(A)** Visual stimulation: six stimuli are presented per second (6 Hz) with a sinusoidal contrast modulation, with oddball stimuli inserted at 6 Hz/5 = 1.2 Hz. **(B)** Stimulation sequences: base stimuli consisted of pseudo-fonts (coarse level) or pseudo-words (fine level) and the oddball stimuli were either a word or a pseudoword (coarse level) or a regular or irregular word (fine level) appearing every five items. Sequences were repeated 4 times. **(C)** Timeline of a stimulation sequence: each sequence started with a fixation cross (2-5s). Then, the stimulation faded in (2s), reached full contrast and remained for 40 seconds before fading out (2s).

## MATERIAL AND METHODS

### 1. Participants

Children were recruited from French-speaking Belgian schools and speech therapy centers in Belgium. Parents gave written consent for the study, which was approved by the Biomedical Ethics Committee of the Université Catholique de Louvain. One participant dropped out of the study, leaving a final sample of 38 children, among which 14 diagnosed with developmental dyslexia, *M* age = 10.03 years; range = 9.11-11.34 years (5 boys) and 24 typical readers, *M* age = 10.03 years; range = 9.1-12.2 years (12 boys, 3 left-handed). This sample size is similar to those of developmental studies using the same approach (Lochy & Schiltz, 2019: N=17; Crollen et al., 2025; N=12, 2 groups), and a recent comparison of dyslexics vs typical adult readers (Lochy et al., 2025, N=14, 2 groups). All children had normal or corrected-to-normal vision. They were all enrolled in French-speaking schools since preschool. They were all from high SES neighborhood and families (average 3.92 on a 6 points scale, assessed by highest study level of the parents). They were tested in two sessions (behavioral, EEG) during the third trimester of grade 4 or the first trimester of grade 5.

### 2. Behavioral testing

Children were assessed with standardized behavioral tests in three domains (see Table 1): (1) general cognitive functions (non-verbal intelligence (WISC-V; Wechsler, 2005), selective attention (EDA; Willig, Billard, Blanc, Langue & Touzin, 2013), digit span forward and backward (WISC-V; Wechsler, 2005), (2) reading prerequisite skills (Rapid Automatic Naming (RAN) of digits and objects (BALE; Jacquier-Roux, Lequette, Pouget, Valdois, & Zorman, 2010), phonological awareness (BELEC; Mousty & Leybaert, 1999), and (3) reading ability (reading lists of pseudowords, regular and irregular words (BALE; Jacquier-Roux, Lequette, Pouget, Valdois, & Zorman, 2010, LMC-R; Khomsi, 1999) and text reading (Alouette-R; Lefavrais,2005).

**Table 1.**
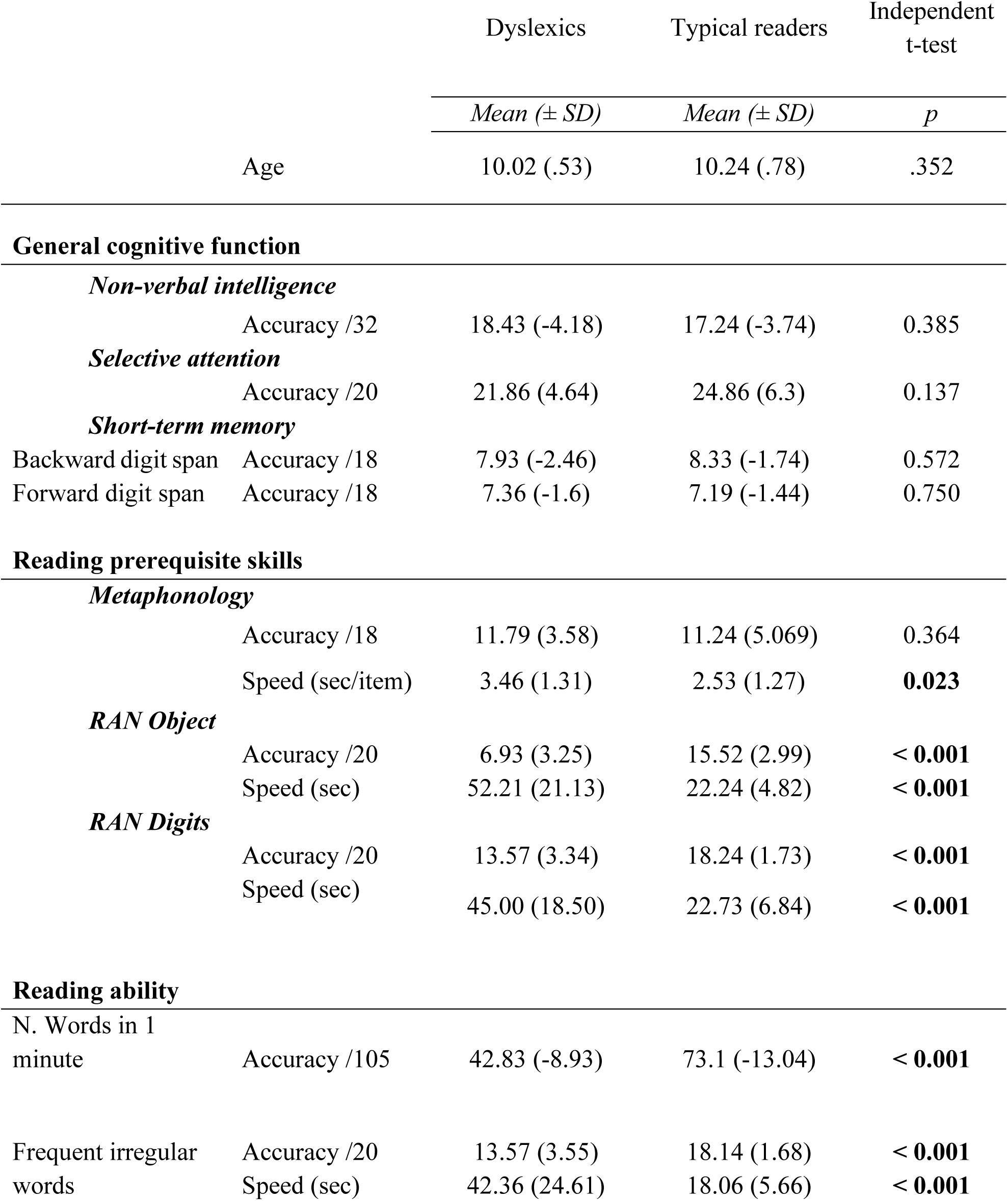
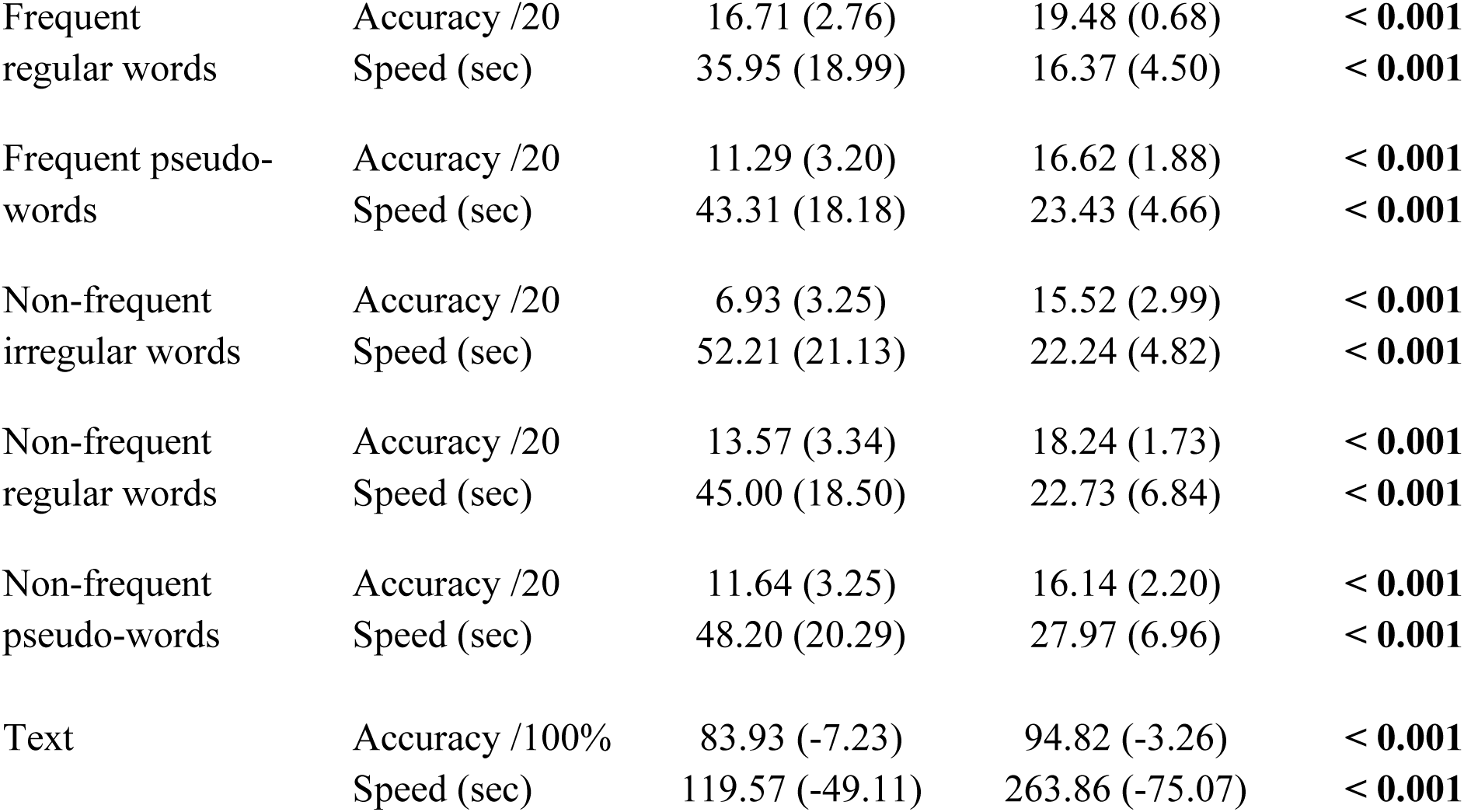
Descriptive statistics of the scores obtained by the dyslexic group (N=14) and the typical readers group (N=21) on behavioral tests.

To identify outliers within the sample distribution, individual Z-scores or standard notes were computed for each test. One typical-reader child was excluded because of scores lower than 2 standard deviations (SD) on non-verbal measures. Results of the behavioral assessment are summarized in **Table 1**. Both groups performed within the norms on general cognitive functions, but children with dyslexia differed significantly from their typical reading peers on all reading-related tests.

### 3. EEG testing

#### 3.1. Stimuli

This study included stimuli similar as validated in other studies (van de Walle de Ghelcke et al., 2020, 2021). Letter strings were presented in Verdana font, as well as non-letter string stimuli, ranging from 47 to 77 pixels in height and from 119 to 199 pixels in width. Stimuli ranged from 3.11 to 5.20 (width) and 1.32 to 2.18 (height) degrees of visual angle. Screen resolution was 800 × 600 pixels with a refresh rate of 60 Hz. Four experimental conditions were designed: two of them implying coarse discrimination level and two others a fine-grained lexical discrimination.

At a coarse level of discrimination (Fig.1B.), pseudo-font strings (PF, N = 20) constituted the base stimuli. In the first condition, they were interspersed with words (W, N = 20), and in the second condition, with pseudo-words (PW, N = 20). Stimuli were built in several steps. First, words of four (N = 10) or five letters (N = 10) were selected from the Manulex database (Lété et al., 2004). In a second step, one pseudoword was built from each word by changing the position of their constitutive letters (e.g. the words ‘fleur’ and ‘neige’ give rise to the pseudo-words ‘frelu’ and ‘igene’). All pseudo-words were pronounceable letter strings respecting the phonological rules in French. The number of letters, letter identities, and bigram frequency were matched between words and pseudo-words (t(19)= .427; p = .675; PW = 8,141.15 mean ± 780.70 SD, W = 8,390.10 mean ± 952.97 SD). The base pseudo-font strings were built from the words and pseudowords (**Fig. 1B**) by applying a vertical flip and segmenting their letters into basic features with Adobe Photoshop. Segments were rearranged to create pseudo-letters respecting the original word size and number of characters. The pseudo-letters included junctions, ascending or descending features, and close-up shapes as real letters. Therefore, each oddball stimulus (W, PW) had a corresponding pseudo-font stimulus with the same black-on-white contrast, ensuring comparability in terms of low-level visual properties.

For the fine-grained discrimination level (Fig1.B), two other experimental conditions were designed by embedding regular (REG, N = 20) or irregular (IRR, N = 20) words into base pseudoword strings (PW, N = 20). We ensured that regular and irregular words had the same number of letters by choosing words of five (N = 10) or six letters (N = 10) for each category.

Also, they were matched on bigram frequency (t(19) = -.114; p = .910; REG = 12,076.50 mean ± 3,200.982 SD, IRR = 12,292.35 mean ± 7,733.29 SD) and orthographic neighborhood (t(19) = .705; p = .490; REG = 1.35 mean ± 1.95 SD, IRR = 0.90 mean ± 1.99 SD). Once again, a pseudoword was built for each regular or irregular word by changing the positions of its constitutive letters. Regular words and their matched pseudowords did not differ in bigram frequency (t(19) = 1.03; p = .315; W = 1.35 mean ± 1.95 SD, PW = 1.05 mean ± 1.05 SD), or orthographic neighborhood (t(19) = .562; p = .581; REG = 12 076.5 mean ± 3 200.9 SD, PW = 11 094.5 mean ± 4 625.9 SD). Similarly, irregular stimuli adhered to these criteria, showing matched bigram frequency (t(19) = .285; p = .779; W = 12 292.3 mean ± 6 733.2 SD, PW = 11 845.7 mean ± 3 046.1 SD) and orthographic neighborhood (t(19) = -.515; p = .612; W = 0.90 mean ± 2 SD, PW = 1.10 mean ± 1.55 SD).

#### 3.2. Procedure

The child sat 1 m away from the computer screen in a quiet room at school or in the laboratory. A fixation cross appeared in the center of the screen on a gray background 2-5 s before the stimulus sequence. Stimulation sequences lasted 40 seconds flanked by a 2 second fade-in and fade-out period to avoid abrupt eye movements or blinks.

The procedure was similar to previous FPVS-EEG studies with children (Lochy et al., 2016; Lutz et al., 2024; van de Walle de Ghelcke et al., 2020). Regardless of conditions, every sequence had the same structure: the oddball or deviant (D) stimuli appeared every fifth stimuli, such as: BBBB*D*BBBB*D*BB… Note that responses amplitudes are not affected by item repetition rate (the fact that base stimuli are repeated more often than deviant stimuli) in the FPVS-oddball paradigm, at least with large sets of items (Lochy et al., 2024; Retter & Rossion, 2016; Retter et al., 2020). Stimuli were randomly presented with no immediate repetition through sinusoidal contrast stimulation with Java SE Version 8. We displayed the base stimuli at a frequency of 6 Hz, resulting in a stimulation rate of 1.2 Hz (F/5) for the oddball stimuli. However, for three dyslexic participants, stimuli from the base category were inadvertently presented at 7.5 Hz, which led stimuli from the oddball category to occur at 1.5 Hz. Their data could nevertheless be included (see section *Frequency-domain analysis*), and supplementary material presents the individual amplitude responses at the base (**Table 3**) and oddball (**Table 4**) frequencies, showing that responses of these 3 individuals are in the range of those stimulated at 6Hz. Children watched four times each condition in which oddball stimuli appeared 48 times over 40 seconds: words in pseudo-fonts (PF-W), pseudowords in pseudo-fonts (PF-PW) at a coarse discrimination level, and regular words in pseudowords (PW-REGW), irregular words in pseudo-words (PW-IRRW) at a fine discrimination level. Six additional conditions were tested for a related project and order was randomized across participants. Each sequence was started manually to ensure low-artifact EEG signals, allowing a pause of approximately 30 seconds. The total stimulation time amounted to 25 minutes. Participants’ attention was maintained by asking them to press the space bar with their preferred hand when the fixation cross changed from blue to red.

#### 3.3. EEG acquisition and preprocessing

The signal was acquired at 512 Hz using a 32-channels Biosemi Active II system (Biosemi, Amsterdam, Netherlands) with standard 10–20 system locations, plus a row of posterior electrodes including PO9, I1, Iz, I2, PO10 for a total of 37 channels. Electrodes offsets were held below 50 Mv.

Data were processed in Letswave 6 on Matlab in accordance with the methodology employed in similar studies (Lochy et al., 2024; Retter & Rossion, 2016). A *Butterworth* filter (range = 0.05-100Hz) and a *Multinotch* filter (at 50Hz and 100Hz, width = 0.5 Hz, slope = 0.5Hz) were applied to remove electrical noise. EEG data were segmented into 46-second sequences, including fade-in and fade-out periods. Eye blinks and movements were removed in an independent component analysis (ICA). Noisy channels (less than 2% in total) were identified through visual inspection and interpolated with neighboring channels equally across conditions. One sequence was removed for five participants and two to five sequences were removed for four participants, ensuring that at least 3 repetitions of the same condition remained in the data for further averaging. All channels were referenced to a common average channel. Then, EEG data were re-segmented to include only the 40 s period of stimulation. Resulting sequences were averaged per participant and condition in the time-domain to increase signal-to-noise ratio (SNR).

#### 3.4. Frequency-domain analysis

A Fast Fourier Transform (FFT) was applied and a normalized amplitude spectrum (0-256 Hz) was extracted for all channels. The FFT segments were chunked into epochs to isolate responses at the oddball frequency and harmonics, and to combine them in further analysis. The chunking procedure is detailed in Figure 2 for the base-6Hz (*f*) stimulation, and in supplemental material for base-7.5Hz (*f*) stimulation **(SuppMat.1)**. The duration of each epoch remained 1.124 Hz in both cases, which enables the alignment of the chunks, in which the response at the oddball frequency (or harmonic) is centered (**Fig2.** and **SuppMat1**). The interval was contingent to the frequency of the oddball stimuli: 1.2 or 1.5 Hz (f/5). Each chunk included 25 bins, with the bin corresponding to the frequency of category change in the middle (the 13^th^) and 12 bins on either side. A range of 10 bins on each side of the frequency bin of interest was used to define the baseline brain’s electrical activity, with adjacent bins and extreme amplitudes excluded. To quantify the EEG response in microvolts (μV), the baseline brain’s electrical activity was subtracted from the amplitude response at the frequency of interest (such baseline corrected amplitudes are displayed in **Figure 2B**, while the spectra in **Figure 2A** shows raw FFT amplitudes). A grand average was calculated for each condition and group prior to determining the significance of the EEG response at the oddball frequency and harmonics, as well as at the base rate and harmonics. Subsequently, z-scores were calculated as follow **Z(x) = [x-baseline mean] / [baseline standard deviation]** and computed at every channel. The significant baseline corrected amplitudes (p < .05, one-tailed, signal > noise) at frequencies of interest and their harmonics were summed regardless of group or condition in order to quantify the periodic response distributed over multiple harmonics (Retter et al., 2021) (in Fig2.B, this sum is represented by the red line).

**Fig 2.**
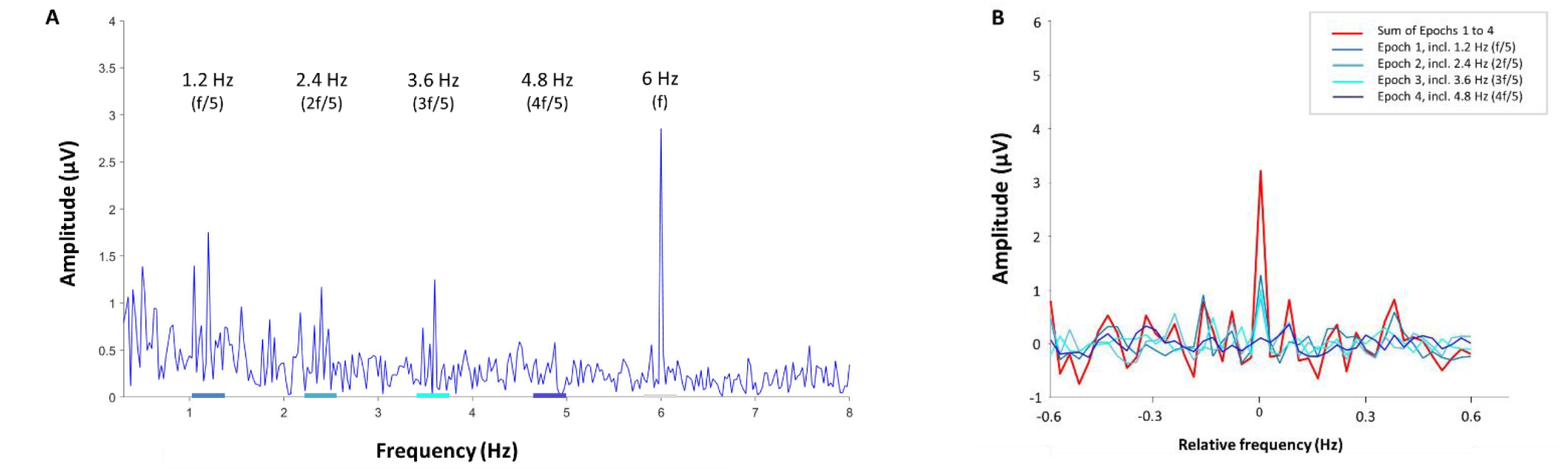
Example of FFT spectra at 6Hz, and its segmentation into chunks allowing for a combination of responses (see also SuppMat 1). A: The FFT spectrum of a 6 Hz stimulation signal and its harmonics in one single participant for the coarse condition PF-W. The base rate response is clearly visible at 6Hz, with a grey shade, and the oddball responses at 1.2Hz (and harmonics), with blue shades on the X-axis indicating the portion of the FFT segment that is being “chunked” for further processing. Segmentation started at 0.637 Hz with a duration of 1.124 Hz. B: The superimposed segmented chunks (shaded blue) and the sum (red) of the baseline corrected FFT segments or “chunks” containing the response of interest centered and surrounded by 12 bins on each side (10 bins were used to compute the baseline correction).

### 4. Principal Component Analysis of the behavioral data

A composite score was computed by calculating the ratio of accuracy to time (in seconds) for each child and each reading subtest. The scores for digits span (backward, forward), the non-verbal intelligence and the selective attention tasks were retained as input variables for Z-scores standardization as these tests are not timed. Principal Component Analysis (PCA) was then used to analyze and reduce the number of variables. A total of 14 variables with *variance ≠ 0* were retained for analysis. All variables were scaled and centered (Z-scores) based on the entire sample. The suitability of the data for PCA was confirmed by a Kaiser-Meyer-Olkin (KMO) value exceeding 0.5 and a significant Bartlett’s test of sphericity (p < 0.05). Principal components were extracted based on eigenvalues exceeding 1. A Varimax rotation was applied with the objective of maximizing the correlation of variables within components while ensuring that each principal component provided distinct information. Loadings with absolute values exceeding 0.5 were retained for interpretation. The highest factor loading was used when variables exhibited cross-loadings across multiple components.

## RESULTS

Analyses were performed on 36 children after excluding the data of two children (one child with dyslexia and one typical reader) because of amplitudes responses above 2.5 SD in both ROI (left, right) across the conditions. Individual data (in µV) are presented in Supplementary Material (Supp.Mat.2).

### 1. Oddball discrimination responses

As can be seen on Figure 3 (coarse-grained discrimination) and Figure 4 (fine-grained lexical discrimination), clear responses occurred for oddball stimuli, in both groups, and displayed a left temporo-parietal topography. These responses at 1.2Hz were significant (z-scores > 1.64) up to seven harmonics (from 1.2Hz to 8.4Hz, excluding the base rate 6Hz) on a left parieto-temporal region including electrodes PO9 and P7 (Left ROI). A contralateral ROI with electrodes PO10 and P8 was defined for hemispheric comparison. Individual data (in µV) for children with dyslexia are presented in Supplementary Material (Supp.Mat.3).

**Fig 3.**
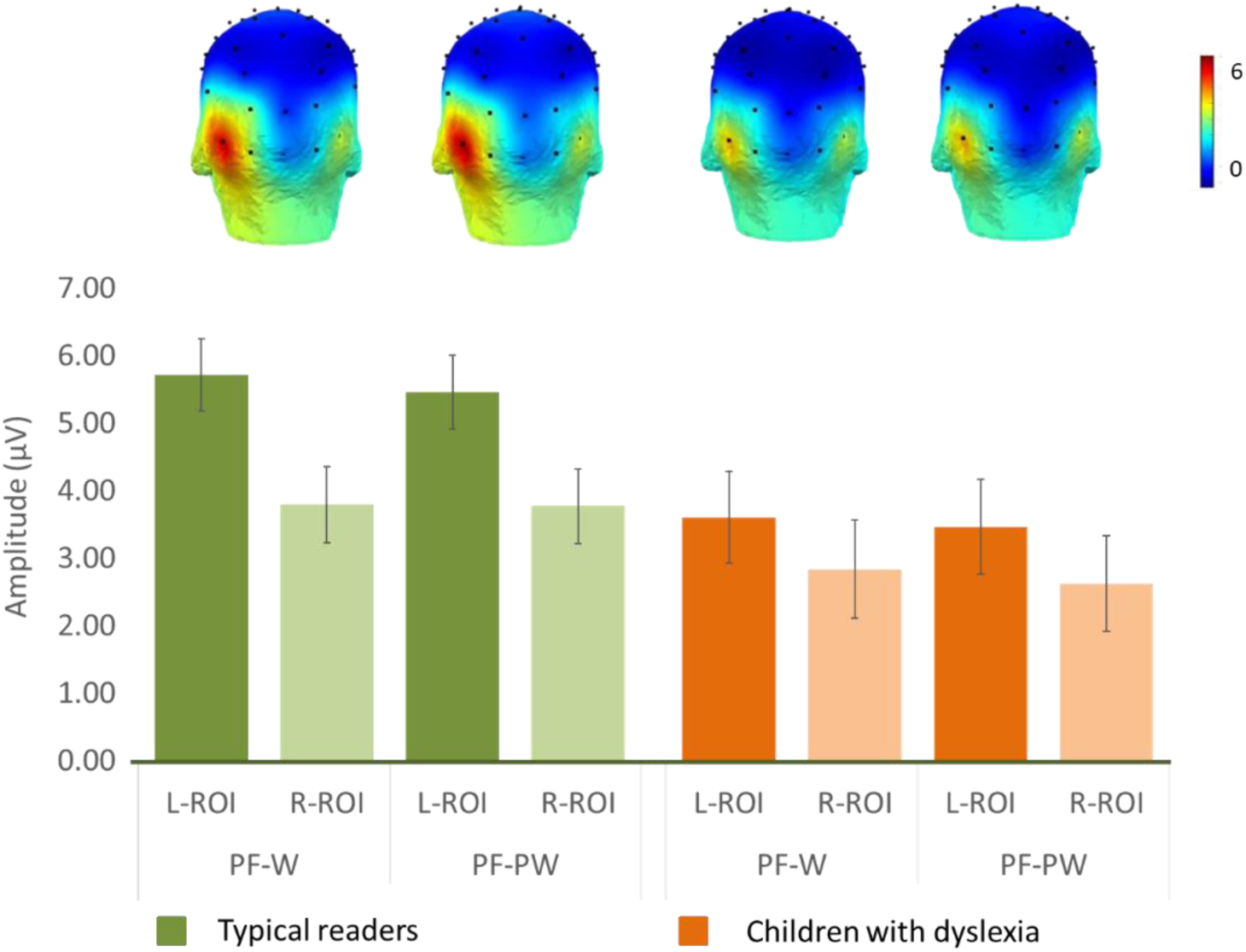
Print-selective responses during coarse-grain discrimination of words (W) or pseudowords (PW) in pseudo-fonts (PF). EEG amplitudes (in μV, sum of harmonics) are shown by Group (*green:* typical readers; *orange:* children with dyslexia) in the parieto-temporal ROIs. Topographies are shown by group and condition above each corresponding bar.

**Fig 4.**
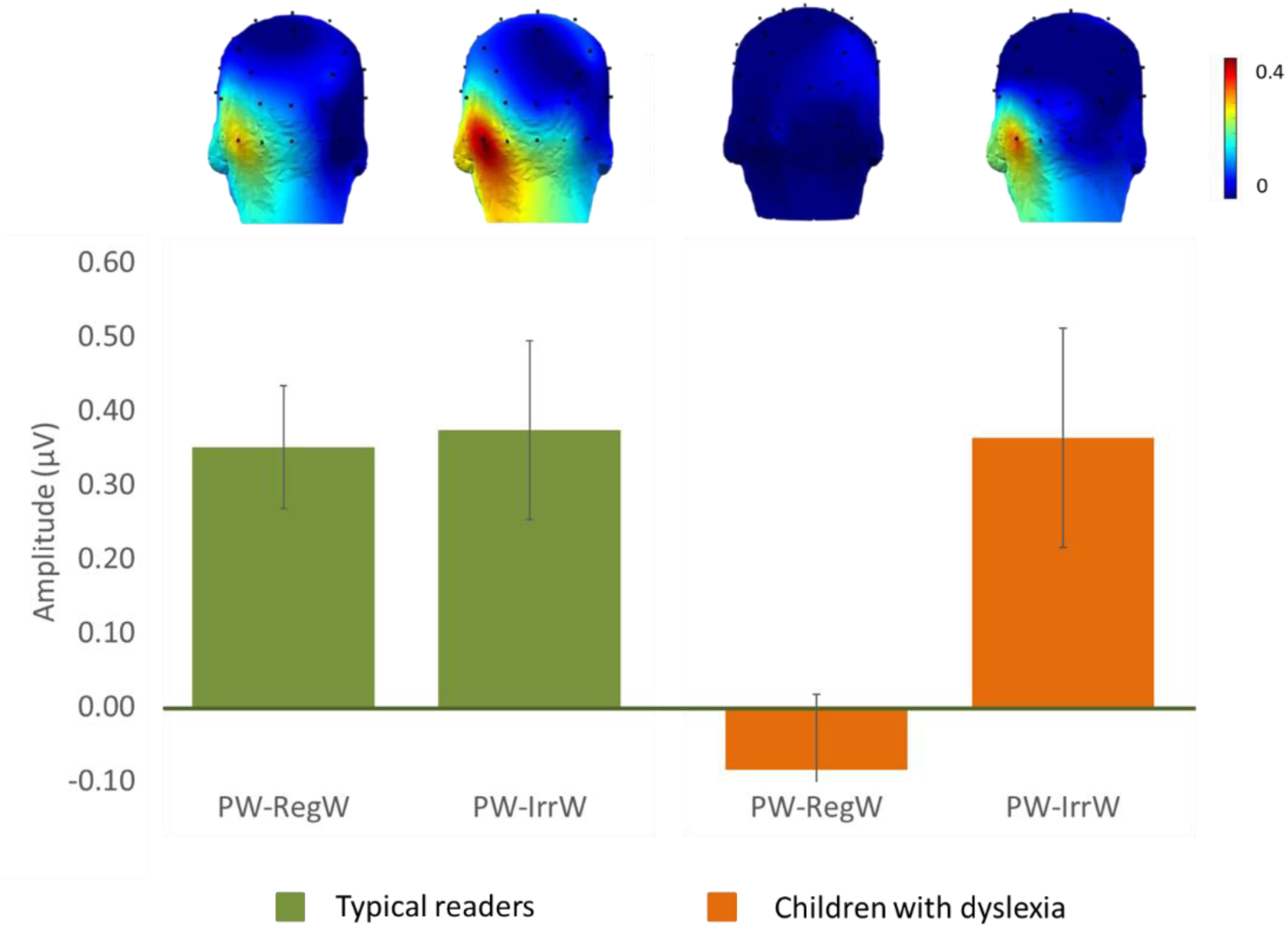
Word-selective EEG amplitudes (in μV, sum of harmonics) in the left ROI during fine-grained lexical discrimination of regular words (Reg) or irregular words (IRR) in pseudowords (PW), shown by Group (*green*: typical readers; *orange*: children with dyslexia)

#### 1.1. Coarse-grained level of discrimination: print among pseudofonts

The sum of baseline-corrected amplitudes for the seven significant harmonics at the oddball frequency in the parieto-temporal ROIs were submitted to an ANOVA with *Conditions* (PF-W, PF-PW) and *ROI* (Left vs. Right) as within-subjects factors, and *Groups* (Dyslexics vs. Typical-readers) as a between-subjects factor. As illustrated in **Figure 3**, we observed a significant main effect of ROI, *F*(1,33) = 15.04; *p*<.001; partial *η2* = .313, reflecting larger amplitudes in LROI, *M*= 4.36µV; *SD* = .293, than in RROI, M= 3.01µV; *SD*= .295. Concerning the main effect of *Group*, although letter-selective responses were almost twice larger in typical readers (3.30 µV) than in dyslexics (1.81 µV), the difference was only marginally significant *F*(1,33)= 3.55; *p =* .06; partial *η2* = .097. There was no main effect of *Conditions F*(1,33)= 1.507; *p =* .228, response amplitudes being equivalent for discrimination of words (PF-W condition: 3.98µV) or pseudo-words (PF-PW condition: 3.82µV). The interactions were not significant: *Hemisphere*Group F*(1,33)= 1.507; *p =* .228, *Condition*Hemisphere F*(1,33)= .397; *p =* .533, and *Condition*Hemisphere*Group F*(1,33)= .560; *p =* .460. (see Figure 3)

#### 1.2 Fine-grained lexical discrimination: words among pseudo-words

A visual inspection of scalp topographies (**Figure 4**) and z-score analysis per harmonic (see Method) suggested no response in the right hemisphere, therefore we performed a one sample t-test to examine if the response amplitudes in the right ROI differed significantly from 0. The results disclosed that they were not different from 0 for both regular words (PW-RegW) (t(34) = 1.39, *p =* .17) and irregular words among pseudowords (PW-IrrW) (t(34) = -.50, *p =* .61).

Therefore, we focused the analysis on the left hemisphere and conducted a repeated measure ANOVA on the sum of baseline-subtracted amplitudes responses to words with *Conditions* (PW-RegW vs. PW-IrrW) as within-subjects factor and *Groups* (Dyslexics vs. Typical readers) as a between-subjects factor. This analysis showed a significant main effect of *Conditions*, *F*(1,33) = 8.85, *p =* .05, *partial η2* = .211 : within matched pseudowords strings, irregular words induced higher response amplitudes than regular words (respectively .43 µV and .12 µV). No main effect of *Group* was observed *F*(1,31) = 2.403; *p =* .125; *partial η2* = .07, but the interaction *Conditions*Group* reached significance *F*(1,33)= 6.03; *p =* .02; *partial η2* = .155): while irregular words resulted in comparable amplitudes responses in both dyslexic (.445 µV) and typical readers (.422 µV), regular words elicited discrimination responses for typical readers (.367 µV) but not for dyslexics (-.124 µV) (**Figure 4**). A t-test for independent samples ensured that the amplitude for regular words was significantly higher for the typical readers than for the dyslexics, *t*(33) = - 3.699; p<.001, while there was no difference between the groups in the PW-IrrW condition, *t*(33) = .112; *p =* .911. Since the response pattern for children with dyslexia in the PW-RegW condition suggested no clear discrimination response for regular words, we ran a one-sample t-test against 0, which showed that amplitudes were not different from 0, *t*(13) = -1.6, *p =* .13.

### 2. Brain-behavior correlations

#### 2.1. Principal Component Analysis of the behavioral data

Four components were extracted (see Table 2). The first one grouped all the composite scores calculated for reading tests (Alouette-R, Bale, LUM): all factor loadings were above .89, and it accounted for more than 53% of variance. We interpreted this component as reflecting “reading ability”. Under the second component, the selective attentional test (EDA), the RAN objects and digits were grouped, all involving visual processing and visuo-motor coordination, cognitive processing speed, as well as selective and sustained attention, which could therefore reflect “executive functions”. The third factor included the meta-phonology subtest, digit span and reverse digit span, all tasks requiring holding and manipulating phonological information in working memory and therefore reflect “verbal working memory”. The fourth factor indicates “general cognitive ability” with the non-verbal intelligence test (WISC).

**Table 2.**
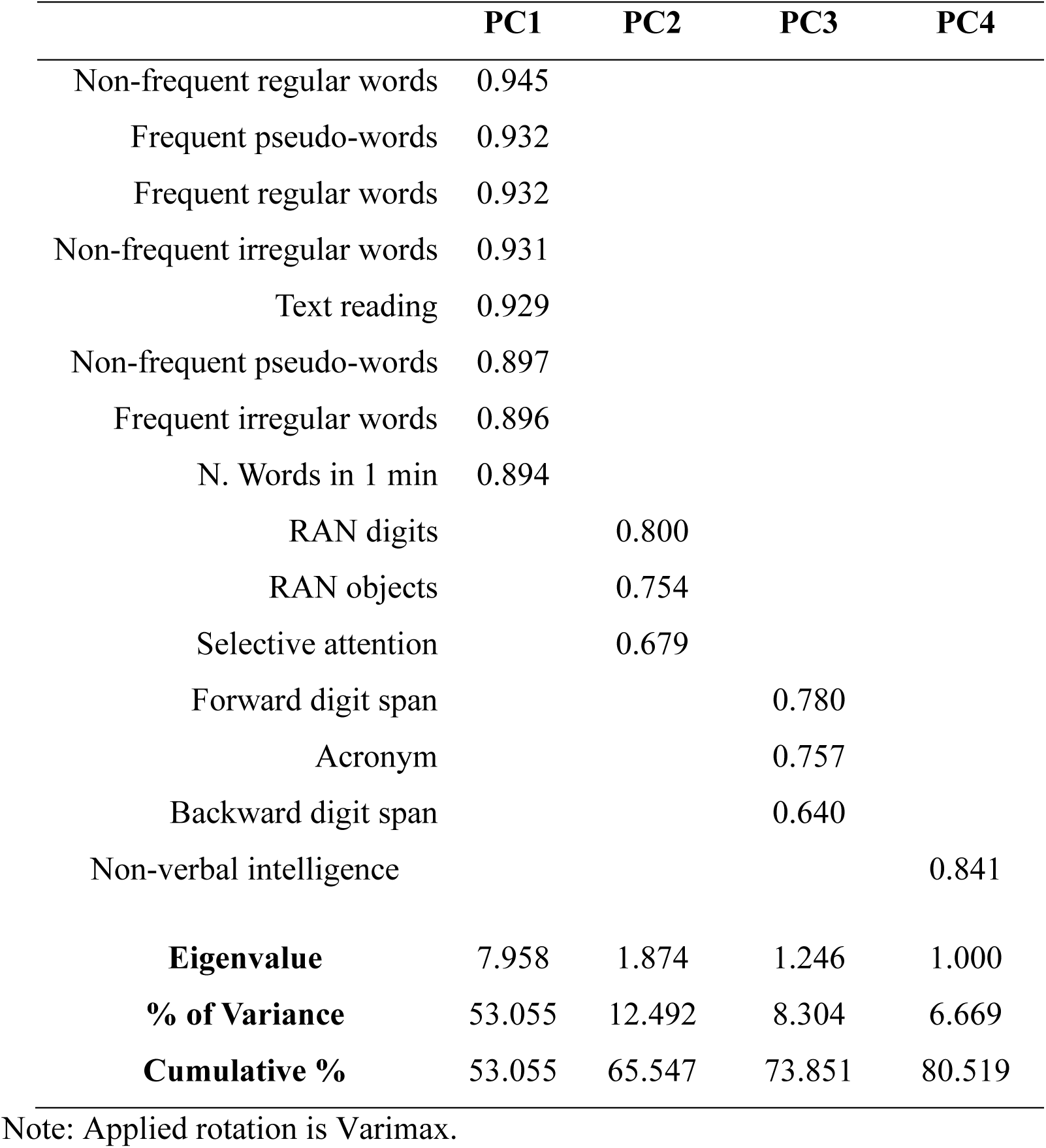
Results of the Principal Component Analysis after varimax rotation of cognitive measures.

|  | **PC1** | **PC2** | **PC3** | **PC4** |
| --- | --- | --- | --- | --- |
| Non-frequent regular words | 0.945 |  |  |  |
| Frequent pseudo-words | 0.932 |  |  |  |
| Frequent regular words | 0.932 |  |  |  |
| Non-frequent irregular words | 0.931 |  |  |  |
| Text reading | 0.929 |  |  |  |
| Non-frequent pseudo-words | 0.897 |  |  |  |
| Frequent irregular words | 0.896 |  |  |  |
| N. Words in 1 min | 0.894 |  |  |  |
| RAN digits |  | 0.800 |  |  |
| RAN objects |  | 0.754 |  |  |
| Selective attention |  | 0.679 |  |  |
| Forward digit span |  |  | 0.780 |  |
| Acronym |  |  | 0.757 |  |
| Backward digit span |  |  | 0.640 |  |
| Non-verbal intelligence |  |  |  | 0.841 |
| **Eigenvalue** | 7.958 | 1.874 | 1.246 | 1.000 |
| **% of Variance** | 53.055 | 12.492 | 8.304 | 6.669 |
| **Cumulative %** | 53.055 | 65.547 | 73.851 | 80.519 |

Note: Applied rotation is Varimax.

Thus, only the first component related to reading abilities, all others reflecting rather general cognitive functions. The current analysis therefore focuses on the first component only: individual scores were derived for this first component and correlated with EEG amplitudes to investigate the relation between behavioral measures of reading ability and the neural responses to words measured by EEG.

#### 2.2. Relation between reading ability and neural responses

Given that the non-significance of the group effect at a coarse level of discrimination warrants further consideration, Spearman correlations (one-tailed) were computed between the first component of the PCA reflecting reading abilities, and the left ROI described above (**Figure 5**). At a coarse-grained level of discrimination, a significant and strong positive correlation was found between participants’ reading performance (PC1) and the oddball responses of both words *r_s_*(33) = 0.480; *p =*.002 (Figure 5A) and pseudo-words *r_s_*(33) = 0.459; *p* = .004 (Figure 5B) among pseudofonts. At a fine-grained lexical level, a positive correlation was found between reading ability (PC1) and discrimination of regular words, *r_s_*(33) = .389; *p =* .013 (Figure 5C), but not of irregular words *r_s_*(33) = .021; *p =* .910 among pseudowords. Note that correlations were selective to the oddball responses, highlighting the specificity of the relation between PC1 and neural responses that reflect orthographic and lexical processes. Indeed, no correlation was significant with amplitudes at the base rate (PC1 and coarse-grained responses to words rs(33) = 0.045; p =.800 or pseudowords rs(33) = 0.001; p =.994, and fine-grained responses to regular rs(33) = 0.052; p =.768 or irregular words rs(33) = -0.042; p =.813).

**Fig 5.**
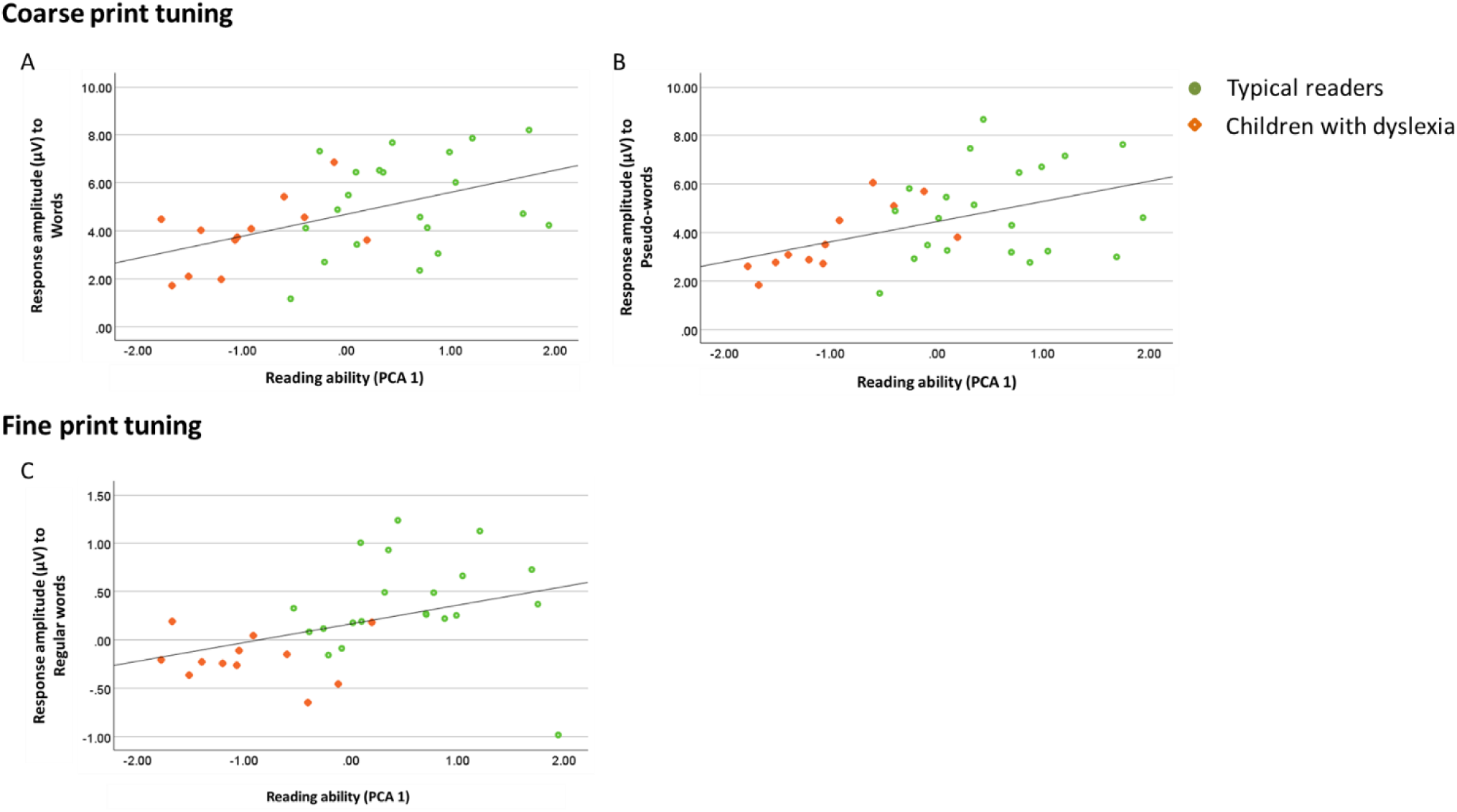
Scatter plots showing the relationship between reading ability (quantified as the first principal component -PCA 1-encompassing reading-related measures) and neural response amplitude (μV) to words in pseudofonts (A) pseudo-words in pseudofonts (B) and regular words in pseudowords (C). Individual data points represent typical readers (*green circles*) and children with dyslexia (*orange diamonds*).

### 3. Base rate responses

To exclude that differences between groups could reflect a difference at a general visual response and attentional level, we examined also responses at the base rate. At the base frequency (6 Hz), the region of interest (ROI) included eight channels (max. 4.62 µV, min. 3.18 µV) in a medial occipital region (see **Figure 6**), as already found in previous studies with the same approach (Lochy et al., 2015; 2016). Responses were significant (z-scores > 3.1) up to four harmonics (from 6Hz to 24Hz). ANOVAs on baseline subtracted amplitudes at the occipital medial ROI were conducted separately for coarse and fine-grained contrast levels. The within-subjects factor was *Conditions* (coarse-grained level : PF-W vs. PF-PW; fine-grained level : PW-REGW vs. PW-IRRW) and the between-subjects factor was *Groups* (Dyslexics, Typical readers). Both analyses showed no main effects or interactions. For the coarse contrast: *Conditions F*(1,33) = 1.214; *p*=.27; *Groups F*(1,33) = .013; *p=*.954 and interaction *F*(1,33) = 2.438; *p=*.128 (**Figure 6A**), and for the fine contrast : *Conditions F*(1,33) = 1.234; *p*=.275; *Groups F*(1,33) = .378; *p=*.802; and interaction *F*(1,33) = .378; *p=*.543 (**Figure 6B**).

**Fig 6.**
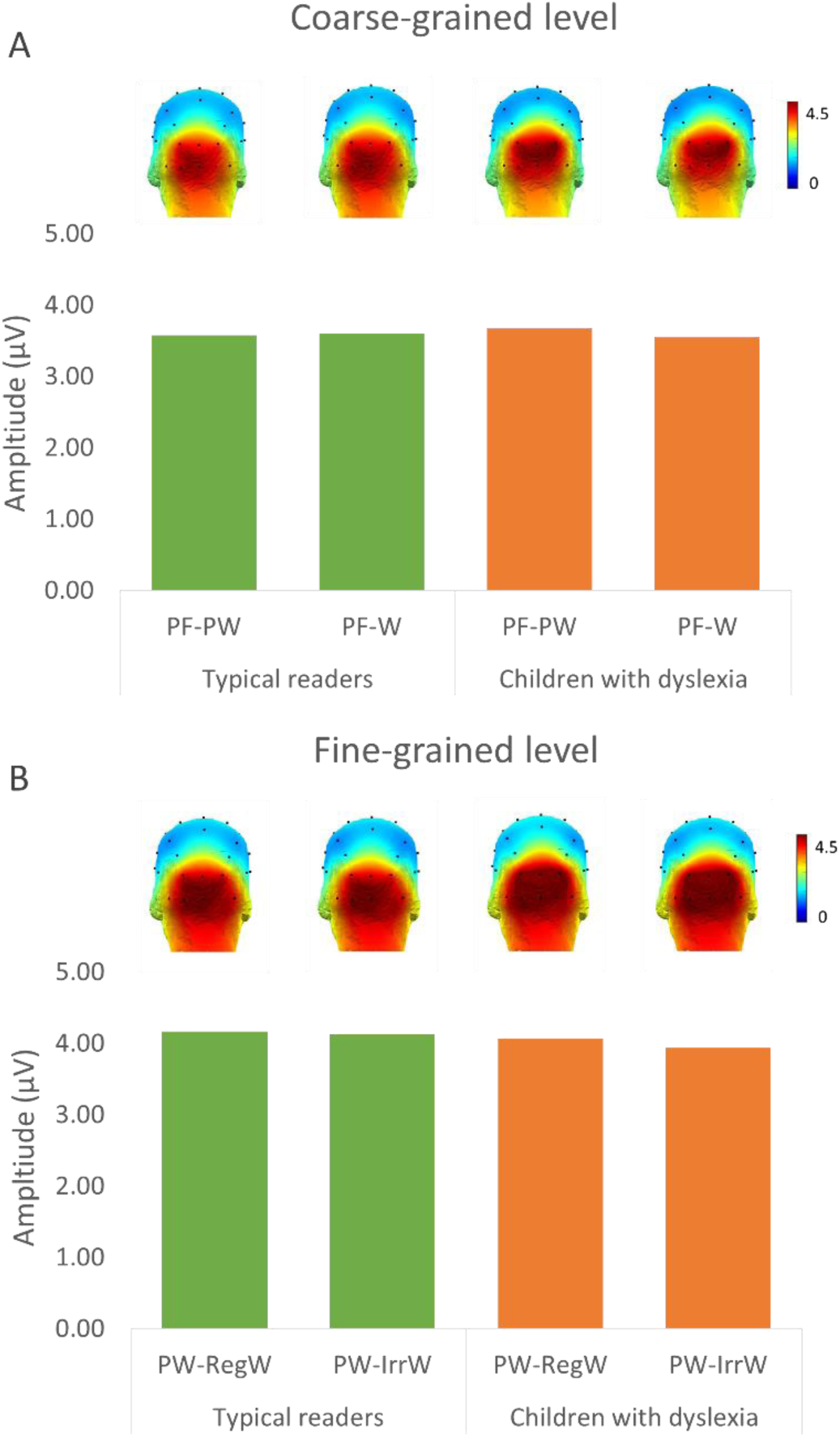
Base rate response amplitudes (in μV) at the medial-occipital region of interest during **A.** coarse-grain processing (Pseudo-words (PW) or Words (W) in pseudofonts (PF) and **B**. fine-grain processing (Regular words (Reg) or Irregular words (IRR) in pseudo-words (PW), shown by Group (*green*: typical readers; *orange*: children with dyslexia)

## DISCUSSION

Neural responses for visual word recognition were investigated in 10-year-old typical readers and age matched children with dyslexia through FPVS-oddball design with EEG recordings. Two levels of contrasts were implemented: a coarse-grained level, which assesses print sensitivity by displaying letter-strings oddball stimuli among non-letter strings, and a fine-grained level, which assesses lexical recognition by displaying words among pseudowords. Overall, discrimination responses for the oddball stimuli were located over a left occipito-temporal site on the scalp at the exact frequency of category change and its specific harmonics. In the coarse-grained contrast, words and pseudo-words were equally discriminated from pseudo-fonts in each group, and children with dyslexia exhibited non-significantly lower amplitudes than typical readers. A continuous variable approach, revealed that behavioral reading abilities were strongly related (rho=0.48) with print-selective response amplitudes. In the fine-grained contrast, irregular words were distinguished from pseudo-words similarly in children with dyslexia and typical readers, whereas regular words in pseudowords strings elicited no discrimination at all in children with dyslexia. Overall, the FPVS-EEG is a robust novel approach to address the developmental origin of sensitivity to lexicality that sheds light on differences between ten-year-old dyslexics and typical readers. This approach has the advantage of obtaining results very quickly (4 repetitions of 40 s per condition), with neural responses reflecting automatic word processing via an implicit task (children do not have to read or even explicitly process the written stimuli). These aspects make it suitable for testing young and clinical populations.

### Lexical responses in 10-year-old children

The developmental trajectory of fine-tuning sensitivity to word forms has not yet been clearly disclosed. ERP studies suggest that it may emerge between 9 and 11 years of age, but the findings are inconsistent. The emergence of automatic recognition that letter strings are words rather than pseudowords is an important marker of rapid lexical access. The present study is the first to reveal clear lexical responses among10-year-old children, both typical and dyslexic readers, using an FPVS-oddball design with words periodically embedded in pseudowords at 6Hz (166ms per stimulus). Previous FPVS-oddball designs have reported discrimination between words and pseudowords in adults (Lochy et al., 2015, 2024; Marchive et al., 2025) but not in first, second and third graders (Lochy et al., 2016; Van de Walle de Ghelcke et al., 2020; Lutz et al., 2024). To date, only one FPVS-EEG study (Wang et al., 2023) has reported lexical (words within pseudowords) and sublexical (pseudowords within nonwords) responses in seven-year-old proficient readers. These authors implemented an alternative frequency-tagging paradigm that differs from the oddball design used here in terms of task demands and periodicity characteristics. In their study, three-letter English stimuli were displayed at a base frequency of 2 Hz (every 500 ms) with a word stimulus appearing at 1 Hz (every 1000 ms), so that a word systematically alternated with a pseudoword. Participants had to press a button when a stimulus was repeated three times in a row, a task requiring to explicitly focus on the linguistic stimuli being displayed. Thus, at this slow stimulation rate, it is possible that the EEG response reflects both phonological decoding and orthographic processing. This differs from the present design, in which the short duration of stimuli (166 ms) emphasizes rapid automatic lexical access from the orthographic form. In addition, the unrelated color change detection task is strengthening attention without any instruction related to the linguistic aspects of stimuli. Nevertheless, future studies should examine if reduction in the presentation rate in the oddball paradigm may enable further insights into different aspects of lexical access.

### Automatic visual word recognition is modulated by discrimination level

Lexical responses were investigated at two levels of contrast: firstly, responses to words and pseudowords were compared within pseudo-font base stimuli (coarse-grained level); and secondly, responses to words within pseudowords base stimuli (fine-grained level) were assessed.

At the coarse contrast level, amplitudes at the oddball frequency did not differ between words and pseudo-words contrasted to pseudo-fonts. The same lack of finding was previously described among 5 to 7 years old children (van de Walle de Ghelcke et al., 2020) and 7 to 9 years old (Lutz et al., 2024), suggesting that neural specialization for word processing is not yet discernible until 10 years of age in a rapid implicit processing task (Lutz et al., 2024). However, our data also show that words and pseudo-words can be reliably discriminated from one another at a finer contrast level in 10-year-old, at least in non-dyslexic children. Indeed, even if amplitudes were a lot weaker in that case than in the coarse contrast (around 0.40µV vs around 4µV), statistics clearly demonstrate significant word-discrimination responses in the fine contrast level. This suggests that in the coarse contrast, the discrimination of both words and pseudowords in pseudofont strings relied predominantly on letter recognition (print vs. not print) or orthographic familiarity, rather than lexical characteristics. Therefore, when the contrast between base and oddball stimuli is coarse, the shallow processing fails to elicit specific lexical responses (W>PW). At that coarse contrast level, the recording of significant brain responses over the right occipito-temporal region indeed suggests a latent activation of areas dedicated to visual feature processing, which may have been induced by the visual difference between the letter strings composing (pseudo)words and the pseudo-font strings presented at the base rate. In comparison, fine-grained level oddball responses were exclusively located in the left hemisphere. This result suggests that properties of the base strings constrain the discrimination level of the oddball stimuli. Lexical responses are elicited when higher-level lexical processing is activated by embedding words within pseudowords strings, although it is not obligatorily the case in a coarse contrast. Our findings thus demonstrate that the contrast salience is critical, and that 10-year-old children may have automatic lexical access in a rapid passive viewing task. This result is reminiscent of a similar modulation of discrimination response in children in the domain of face-processing (Lochy, Schiltz & Rossion, 2019). In that case, 5-year-old children watched strings of images displaying faces as oddballs against non-face objects (coarse level), which elicited bilateral face-selective responses, or they watched a stream of faces only, differing in identity (fine-grained contrast). This finer-grained visual discrimination level induced right-lateralized discrimination responses, as typically observed in adults. Thus, it is crucial to carefully consider the level of contrast induced by the FPVS-oddball design to make inferences about the development of specific word- (or face) related neural processes.

### Weaker coarse-grained print tuning in dyslexia

Overall, in our experiment, children with dyslexia tended to show smaller response amplitudes to words presented at the oddball frequency than typical readers. This is consistent with a reduced activation of the left occipitotemporal cortex for written word processing in individuals with dyslexia compared to their typically reading peers (Van der Mark, et al., 2009; Pina Rodrigues et al., 2019; Brem et al., 2020), observed across alphabetic languages (Martin et al., 2016). In the coarse contrast (letter-strings among pseudo-letters), the amplitude difference was substantial between children with dyslexia (1.81µV) and typical readers (3.30µV, i.e., a reduction of 45.15% in dyslexia) but did not reach significance. This may be due to the limited number of participants in the dyslexic group, and the large variability across individuals for both groups. However, a significant positive correlation between reading efficiency measures, which essentially separates the two groups of participants, and EEG oddball response amplitudes was found. Children with dyslexia exhibited the lowest reading scores and corresponding EEG responses, whereas typically developing readers demonstrated higher levels of both reading competency and EEG responses. This aligns with reading skill-dependent tuning for words (Kubota et al., 2018; Brem et al., 2010; Saygin et al., 2016). Importantly, there were no differences in amplitudes at the base frequency of visual stimulation between the two groups, suggesting that both groups did not differ in their overall sensitivity to visual stimulation and attention to these stimuli.

### Impact of word regularity

The left ventral occipitotemporal cortex (VOTC) is key to orthographic processing and exhibits sensitivity to regularity along its posterior-anterior axis (Graves et al., 2010; Brunswick et al., 1999) for regular versus irregular words (Cattinelli et al., 2013; Taylor et al., 2013) and, more generally, for opaque versus transparent orthographies (Paulesu et al., 2001).

Here, the discrimination of French irregular words among pseudowords elicited overall higher amplitudes (.43µV) than French regular words (.12µV), as was recently found also in adults (Lochy et al., 2025) although here it was only clearly the case for children with dyslexia. Notably, in that group, response amplitudes for regular words did not differ from zero, meaning that when sequences of pseudowords were presented at the base rate, regular words presented as oddballs were not processed differently than those pseudowords. This finding thus replicates the same pattern recently observed in French-speaking adults with dyslexia (Lochy et al., 2025). Oddballs were matched to the pseudoword base strings for bigram frequency, orthographic neighborhood and consonant-vowel structure, and both types of words were furthermore matched in lexical frequency. Therefore, modulation of the lexical responses in children with dyslexia can be attributed to orthographic regularity per se.

In consideration of the DRC model (Coltheart et al., 2001), the irregular oddball words can only be processed by the direct lexical route, while for regular words, both lexical and phonological mechanisms can contribute to their recognition. Pseudoword base stimuli, on the other hand, trigger only the indirect phonological route. Therefore, it can be proposed that typical 10-year-old readers processed both types of words with the same lexical processes given that both regular and irregular words gave rise to similar amplitudes of discrimination. To the contrary, in children with dyslexia, regular words did not automatically trigger the lexical route in a phonological context. Impressively, this finding replicates recent data obtained with the same paradigm (but different stimulus sets) in adults with dyslexia (Lochy et al., 2025) and therefore, seems to be a genuine signature of reading impairment in French-speaking individuals. The findings align also with Hoffman and colleagues’ (2015) study, which indicated that the neural network for reading varies based on how readers are balancing direct and indirect pathways. This balance is influenced by reading experience and skill. Our results can furthermore be related to context effects on word reading strategies, as reported behaviorally: presenting words in lists intermixed with pseudowords (vs blocked lists) influence reliance on decoding vs. lexical processing (Castro & Lima, 2010; Kolinsky & Tossonian, 2023), because the presence of pseudowords induce processing of smaller units that are then also applied to words. In the present case, the FPVS paradigm measures a differential response to words, that is influenced by the mechanisms involved in processing the base stimuli. Therefore, once sublexical decoding mechanisms are enhanced by context, children with dyslexia do not switch to lexical processing for regular words, even if they are typically described as displaying a phonological impairment (Ramus, 2014; Hulme et al., 2015). Furthermore, in our current sample, the observed deficit in speed during rapid object naming (RAN) corroborates the idea of a concomitant impairment in lexical access.

The lexical route relies on the (VOTC), the anterior temporal lobe (ATL) and the anterior inferior frontal gyrus (IFG) (Oliver et al., 2017; Schlaggar & McCandliss, 2007), that provide a system for identifying words based on memory (Glezer, Jiang & Riesenhuber, 2009; Kronbichler et al., 2004). Irregular words, given their non-standard spelling-to-sound patterns, are thought to rely relatively more on the activation of orthographic conventions stored in long-term memory (Bowey & Muller, 2005; Cunningham et al., 2002). This facilitates the discrimination of irregular words from pseudowords. Furthermore, the triangle model (Plaut et al., 1999; Harm and Seidenberg, 2004) suggests a semantic contribution proportional to the irregularity of orthography-phonology mappings. At a neural level, increased activation in the anterior temporal lobe, a region linked to semantic aspects of language, has been identified during irregular word reading (Hoffman et al., 2015; Wilson et al., 2012; Provost et al., 2016). Therefore, the observed amplitude responses in both groups for irregular word recognition, as well as for regular words in typical readers, may have been driven by semantic involvement.

Let us note that children with dyslexia showed selective responses similar to typical readers for irregular word recognition in the EEG task, but a broad impairment in reading aloud assessment, highlighting that reading aloud and silent visual word recognition may rely on partially different processes (Barker et al., 1992; Georgiou & Parrila, 2020). Indeed, their reading z-scores were below -2 SD in both accuracy and speed, for all word types of the battery (including irregular words). It is important to consider that the focus of FPVS is on automatic visual word recognition, whereas reading aloud demands the integration of orthographic analysis with phonological output, a process that can be particularly challenging for children with dyslexia (Mahé et al., 2018; Zoccolotti et al., 2018). Nonetheless, irregular words elicited discrimination responses similar to those of typical readers. One possible account is that the reading of irregular words may have been the focus of rehabilitation for children with dyslexia inducing overreliance on whole-word memorization and resulting in orthographic recognition similar between the two groups (Clark et al., 2021).

## Conclusion

The sensitivity of FPVS-EEG disclosed word-selective responses in 10-year-old children with and without reading impairment. Coarse and fine print tuning were investigated by means of oddball responses at coarse (W or PW embedded in PF) and fine (RegW or IrrW embedded in PW) levels of discrimination, highlighting the importance of the contrast’s level in eliciting lexical responses. This approach provides novel important findings on the processing of written words without an explicit task, revealing clear lexical responses in 10-year-old children. Strikingly, a dissociated pattern of regular-irregular word brain activation was found at a fine-grained level for children with dyslexia, replicating recent findings in adults (Lochy et al., 2025). Although further research is needed across a wider range of reading abilities to determine whether this reflects a specific marker of dyslexia, the current study reveals clearly the theoretical (e.g., emergence of automatic lexical processing) and clinical (e.g., assessment of rehabilitation effects) potential implications of FPVS-EEG studies.

## Acknowledgements/ funding

We warmly thank all participants of the study, the schools and the therapists that helped us recruit and test children. This work has been funded by the FNR (Fonds National de la Recherche Luxembourg, C21/SC/16241557/READINGBRAIN). CG is funded by the FNR (FNR-AFR individual grant 18868448). AVDW was funded by the Fund for Human Sciences Research of Belgian National Fund for Scientific Research (FNRS-FRESH 31450987).

## CREDIT statement

Conceptualization: AL, BR, AVDW; methodology: AL, AVDW; investigation: AVDW; Analysis: CG, AL; Writing: CG, AL; reviewing & editing: CS, BR, AVDW; supervision: AL; funding: AL, BR, CS

## DATA AVAILABILITY

All data are available on OpenScienceFramework and will be made public upon acceptance of the manuscript. https://osf.io/q5uej/?view_only=c454e815ad8e4b0ebf57c6cf1012e5ba

## ETHICS APPROVAL STATEMENT

The Ethics Review Panel of the Université Catholique de Louvain has approved this study that conforms to the Declaration of Helsinki.

