## Supplementary material for "Lexical brain responses in 10-year-old children are impaired in dyslexia: an FPVS-EEG study"

### Supplementary material associated with the article entitled :
#### "Lexical brain responses in 10-year-old children are impaired in dyslexia: an FPVS-EEG study"

**Supplementary material 1.** Alignment of chunks besides different stimulation rates at 6Hz and 7.5Hz

The Figure below shows in two participants the whole response spectra for responses in the frequency-domain with stimulation at 6Hz (oddball responses at 1.2Hz, 2.4Hz, etc) and stimulation at 7.5Hz (oddball responses at 1.5Hz, 3Hz, etc.). Portions of the segments are chunked around the frequency of interests (highlighted in blue in plots A1 and A2), realigned (plots A2 and B2), and after baseline correction, they are summed (red line in plots A2 and B2).

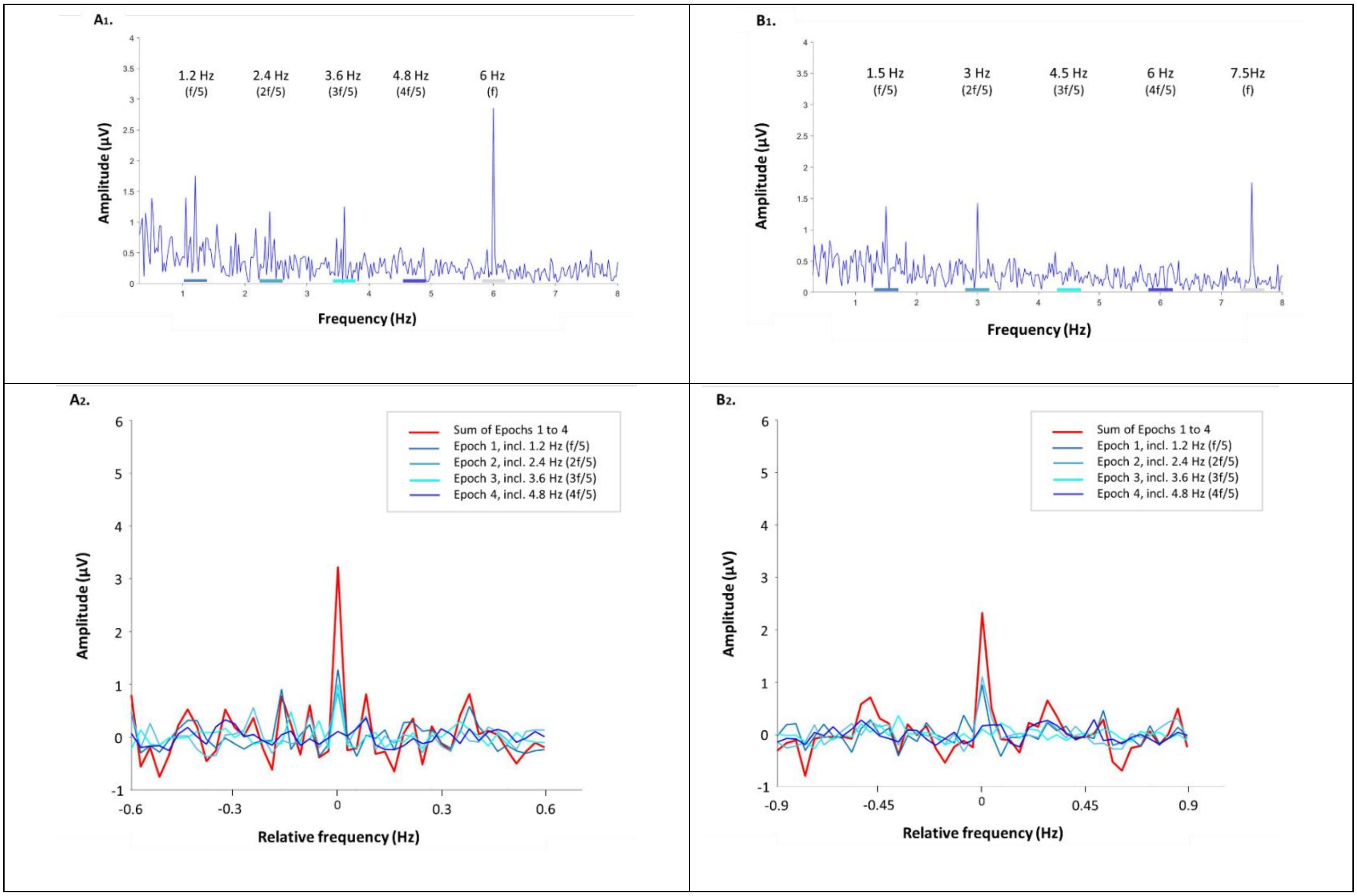

Supp.Fig. 1 **Examples of FFT spectra at 6Hz and 7.5Hz, and their segmentation into chunks allowing for a combination of responses.** A1: The FFT spectrum of a 6 Hz stimulation signal and its harmonics in one single participant for the coarse condition PF-W. B1: FFT spectrum of a single participant with 7.5 Hz stimulation signal and its harmonics, for the same condition. The base rate response is clearly visible at 6Hz/7.5Hz respectively, with a grey shade, and the oddball responses at1.2Hz (and harmonics) and 1.5Hz (and harmonics), with blue shades on the X-axis indicating the portion of the FFT segment that is being "chunked" for further processing. The bottom row (A2-B2 graphs) illustrates the superimposed segmented chunks (or epochs) (shaded blue) and the sum (red) of the baseline corrected FFT segments or "chunks" containing the response of interest centered and surrounded by 12 bins on each side (10 bins were used to compute the baseline correction). The constant duration of each epoch (1.124 Hz) facilitated the alignment of the chunks. This processing approach enables the combination of the discrimination responses despite the use of different stimulation parameters.

**Supplementary Material 2**. Individual EEG Response amplitudes in children with dyslexia: fine and coarse levels at oddball frequency (1.2 Hz vs 1.5 Hz)

| | FINE LEVEL | | COARSE LEVEL | | | |
|---|---|---|---|---|---|---|
| **Subject** | **Regular words** | **Irregular words** | **Words** | | **Pseudo-words** | |
| | | | **RH** | **LH** | **RH** | **LH** |
| S10 | 0.083 | 0.712 | 3.521 | 2.855 | 3.250 | 2.882 |
| S12 | -0.329 | 0.083 | 2.590 | 4.651 | 1.960 | 5.145 |
| S17 | -0.101 | 1.389 | 2.621 | 2.736 | 2.778 | 2.344 |
| S21 | 0.142 | 0.778 | 0.891 | 1.607 | 0.725 | 1.454 |
| S22 | -0.237 | 0.363 | 0.898 | 1.424 | 0.529 | 1.923 |
| S23 | -0.059 | 0.645 | 0.743 | 1.534 | 1.316 | 2.536 |
| S24 | -0.272 | 0.111 | 3.689 | 3.117 | 3.338 | 1.891 |
| S25 | 0.445 | 0.346 | 1.283 | 2.893 | 1.610 | 3.832 |
| S28 | 0.087 | 0.441 | 1.490 | 2.746 | 0.559 | 2.558 |
| S29 | -0.539 | -0.367 | 3.981 | 5.273 | 4.590 | 5.945 |
| S30 | -0.052 | -0.045 | 0.913 | 2.594 | 0.654 | 1.685 |
| S31 | 0.197 | -0.124 | 1.497 | 5.137 | 1.284 | 4.298 |
| S37 | 0.276 | -0.251 | 2.218 | 1.350 | 2.202 | 1.478 |
| S26 | 0.197 | 0.184 | 3.848 | 3.676 | 5.952 | 4.508 |
| ***Mean*** | ***-0.012*** | ***0.305*** | ***2.156*** | ***2.971*** | ***2.196*** | ***3.034*** |
| ***Standard deviation*** | ***0.267*** | ***0.470*** | ***1.215*** | ***1.317*** | ***1.631*** | ***1.457*** |

Oddball amplitudes response for each level of visual word processing: fine (regular vs. irregular oddballs in a pseudoword string) and coarse (words vs. pseudo-words oddball in a pseudofont string). At a fine level, response amplitudes are reported only for the left hemisphere, given that they were not significant at the group level in the right hemisphere. At the coarse level, response amplitudes are reported separately for the left (LH) and right (RH) hemispheres. A grey background indicates children stimulated with 1.5 Hz oddball frequency, while others were tested at 1.2Hz. Mean and standard deviation are calculated including the entire group (N=14).

**Supplementary Material 3**. EEG Response amplitudes in children with dyslexia: fine and coarse levels at base rate frequency (6 Hz vs 7.5 Hz)

| | FINE LEVEL | | COARSE LEVEL | |
|---|---|---|---|---|
| **Subject** | **Regular words in pseudo-words** | **Irregular words in pseudo-words** | **Words in pseudo-fonts** | **Pseudo-words in pseudo-fonts** |
| S10 | 6.343 | 5.768 | 5.163 | 4.841 |
| S12 | 4.185 | 4.150 | 2.888 | 3.407 |
| S17 | 5.111 | 5.348 | 4.539 | 4.619 |
| S21 | 2.706 | 2.929 | 2.414 | 2.496 |
| S22 | 2.457 | 1.880 | 2.292 | 2.025 |
| S23 | 3.893 | 3.250 | 3.007 | 3.288 |
| S24 | 4.955 | 4.585 | 4.477 | 4.838 |
| S25 | 3.997 | 4.171 | 2.898 | 3.411 |
| S28 | 6.821 | 7.030 | 6.580 | 6.120 |
| S29 | 2.211 | 2.501 | 2.206 | 2.385 |
| S30 | 5.563 | 5.534 | 5.489 | 5.938 |
| S31 | 8.700 | 8.859 | 6.878 | 6.025 |
| S37 | 1.719 | 1.561 | 1.774 | 1.596 |
| S26 | 2.803 | 2.416 | 2.422 | 2.830 |
| ***Mean*** | ***4.390*** | ***4.284*** | ***3.787*** | ***3.844*** |
| ***Standard deviation*** | ***1.998*** | ***2.080*** | ***1.708*** | ***1.542*** |

Amplitudes response at the base rate for each level of visual word processing: fine (regular vs. irregular oddballs in a pseudoword string) and coarse (words vs. pseudo-words oddball in a pseudofont string). A grey background indicates subjects tested at a base rate frequency of 7.5 Hz, while others were tested at 6 Hz. Mean and standard deviation are calculated including the entire group (N=14).
